# ENNAVIA is an innovative new method which employs neural networks for antiviral and anti-coronavirus activity prediction for therapeutic peptides

**DOI:** 10.1101/2021.03.25.436982

**Authors:** Patrick Brendan Timmons, Chandralal M. Hewage

## Abstract

Viruses represent one of the greatest threats to human health, necessitating the development of new antiviral drug candidates. Antiviral peptides often possess excellent biological activity and a favourable toxicity profile, and therefore represent a promising field of novel antiviral drugs. As the quantity of sequencing data grows annually, the development of an accurate *in silico* method for the prediction of peptide antiviral activities is important. This study leverages advances in deep learning and cheminformatics to produce a novel sequence-based deep neural network classifier for the prediction of antiviral peptide activity. The method out-performs the existent best-in-class, with an external test accuracy of 93.9%, Matthews correlation coefficient of 0.87 and an Area Under the Curve of 0.93 on the dataset of experimentally validated peptide activities. This cutting-edge classifier is available as an online web server at https://research.timmons.eu/ennavia, facilitating *in silico* screening and design of peptide antiviral drugs by the wider research community.

## 1 Introduction

Viruses are an ancient infection agent that replicate inside the cells of living organisms. They are ubiquitous, affecting all species, from bacteria to plants and animals^[1]^, and are incredibly successful due to their genetic diversity, non-uniformity of mode of transmission, efficient replication and capacity for persistence in their hosts^[2–4]^. Viral diseases are difficult to control due to their potential for high pathogenicity, increased resistance to antiviral drugs, continuous evolution of existing viruses and the emergence of novel viruses^[5]^. Viruses are responsible for many human diseases and are the cause of many death annually. Cold sores, influenza, AIDS, and the current COVID-19 pandemic are all caused by viral infection. Zoonotic viruses, such as the Ebola, Zika, West Nile, HIV, SARS-CoV and SARS-CoV-2 viruses, are especially dangerous, as they are not well adapted to the human hosts’ immune systems, and consequently, cause life-threatening diseases. The World Health Organisation estimated in 2017 that influenza alone is responsible for up to 645,832 death annually, and at the time of writing, the COVID-19 pandemic has been responsible for 2,616,000 deaths. Therefore, the development of novel antiviral drugs, including anti-coronavirus drugs, is important to control emerging viral pathogens.

Host defence peptides are ubiquitous elements of the immune system, having been identified in all living species^[6]^. Indeed, the induction pathways of HDPs are highly conserved among the genomes of animal and plant genomes^[6,7]^. Many HDPs have been found to possess antiviral activity. These antiviral peptides (AVPs) are short, typically 8-40 amino acids, cationic and *α*-helical, although AVPs with an overall negative charge and other secondary structures have also been identified^[8]^. Most importantly, AVPs are a promising resource for the development of novel antiviral drugs for the prevention or treatment of viral diseases^[8]^, including those caused by coronaviruses. For example, a subsequence derived from *β*-defensin, P9, possesses potent inhibitory activity against the SARS-CoV and MERS-CoV viruses^[9]^. Other anti-coronavirus peptides include Mucroporin-M1 and HR2P, which inhibit the SARS-CoV and MERS-CoV viruses, respectively^[10,11]^.

This class of potential antiviral agents possesses a number of advantages over conventional non-peptide drugs, as they are highly specific, costeffective to produce while remaining easy to modify and synthesise, and possess a limited susceptibility to drug resistance^[12]^. Although initially AVPs were isolated from plant and animal secretions where they formed part of the host defence mechanism^[13]^, AVPs have also been derived from chemical^[14]^, genetic^[15]^ and recombinant^[16]^ libraries, as well as from rational design^[17]^. AVPs can be divided into two classes based on their mechanism of action: virus-targeting and host-targeting^[18]^. AVPs belonging to the former class focus on the inhibition of viral enzymes involved in transcription and replication^[19,20]^, or the inactivation of viral structural proteins^[21]^. AVPs of the latter class, act as immunomodulators, like interferons^[22,23]^, or target cyclophilins, which are important cellular factors that are hijacked by viruses during replication cycle^[18,24]^. The currently identified AVPs, however, represent only a small subset of a largely unexplored chemical space, with only a few of those being peptide-based antiviral drugs available on the market. Those drugs include Enfuvirtide, the first peptide inhibitor of HIV-1, Boceprevir and Telaprevir, which both act against hepatitis-C^[25]^. A number of databases exist which detail the antiviral activities of AVPs, such as AVPdb^[26]^, DBAASP^[27,28]^, CAMP^[29]^ and APD3^[30]^.

*In silico* methods offer a fast, efficient way of exploring the large chemical space that AVPs inhabit, by minimizing the quantity of peptides that need to be synthesized and experimentally assayed for antiviral activity. A few methods for the prediction of peptide antiviral activity exist, namely AVPpred^[31]^, AntiVPP 1.0 ^[32]^, Meta-iAVP^[33]^, Firm-AVP^[34]^ and the method of Chang et al.^[35]^. Antiviral peptide prediction methods have been comprehensively reviewed by Charoenkwan et al.^[36]^. Furthermore, Pang et al. recently developed a novel method for the prediction of peptides with specifically anti-coronavirus activity^[37]^. The most popular machine learning methods employed are support vector machines or random forests, although a number of others have also been trialled. Many areas of bioinformatics have benefited from the predictive power of deep learning; neural network-based methods exist for many tasks, such as DeepPPISP for the prediction of protein-protein interaction sites^[38]^, SCLpred and SCLpred-EMS for protein subcellular localization prediction^[39,40]^, CPPpred for the prediction of cell-penetrating peptides^[41]^, HAPPENN for the prediction of peptide hemolytic activity^[42]^, ENNAACT for the prediction of peptide anticancer activity,^[43]^ and APPTEST for the prediction of peptide tertiary structure^[44]^. As the quantity of antiviral peptide sequence data continuously increases, we have exploited the available data to create a deep neural network method for the identification of antiviral peptides from the primary sequence. Herein, we describe ENNAVIA, a novel neural network peptide antiviral and anti-coronavirus activity predictor. ENNAVIA is available as a free-to-use online webserver for the benefit of the academic community at https://research.timmons.eu/ennavia.

## 2 Methods

### 2.1 Datasets

To facilitate easy comparison with existing peptide antiviral activity predictors, the two AVPpred datasets of Thakur et al. were used in this work^[31]^. The first dataset consists of 604 peptides with experimentally validated antiviral activities, and 452 peptides that were experimentally found to have poor or no antiviral activity. This dataset is divided into training and external validation subsets, termed T^544p+407n^ and V^60p+45n^ respectively, where *p* and *n* denote the number of positive and negative samples. For brevity, these are collectively referred to as ENNAVIA-A. The second dataset consists of 604 peptides with experimentally validated antiviral activities, and 604 negative peptides from the AntiBP2 negative dataset, which were randomly extracted from non-secretory proteins^[45]^. This second dataset is similarly divided into training and external validation subsets, termed T^544p+544n^ and V^60p+60n^ respectively, where *p* and *n* again denote the number of positive and negative samples. These are collectively referred to as ENNAVIA-B. Peptide sequences in the datasets consist only of natural amino acids; peptides that contain residues not included in the canonical 20 amino acids are excluded, as are peptides with a sequence length shorter than 7 or longer than 40. Information about the peptides’ secondary structure is not included in the dataset. The datasets are available for download from the webserver website and as supplementary material to this article.

In order to develop a classifier specific to the prediction of peptides with anti-coronavirus activity, two additional datasets were created, ENNAVIA-C and ENNAVIA-D. The positive samples of both datasets are peptide sequences with anti-coronavirus activity, taken from the dataset created by Pang et al.^[37]^. The original dataset included 139 peptide sequences with anti-coronavirus activity. Once peptide sequences with a sequence length shorter than 7 or longer than 40 were excluded, 109 peptide sequences remained. The negative samples of ENNAVIA-C and ENNAVIA-D are the same as the negative samples of ENNAVIA-A and ENNAVIA-B, respectively.

### 2.2 Model validation

It is imperative to thoroughly validate classifier models created by machine learning. Tenfold cross-validations and validation by an external test set were employed for the performance evaluation of all models presented herein. The models trained under cross-validation were ensembled and evaluated with the external test sets. For ENNAVIA-A and ENNAVIA-B, the peptides used in the external test sets are those from the V^60p+45n^ and V^60p+60n^ datasets of Thakur et al.^[31]^, in order to facilitate a direct comparison with existing methods.

Peptides with anti-coronavirus activity which are also present in the ENNAVIA-A and ENNAVIA-B datasets are assigned to the same fold as in ENNAVIA-A and ENNAVIA-B. In order to prevent overfitting, the CDHIT-2D program^[46,47]^ was used to identify anti-coronavirus peptides that can be matched to anti-virus peptides using a sequence identity cut-off value of 0.9. Anti-coronavirus peptides which had high sequence identity to antivirus peptides in the ENNAVIA-A and ENNAVIA-B datasets were assigned to the same fold as those peptides. The negative peptides of the ENNAVIA-C and ENNAVIA-D datasets maintained the same fold-assignment as in the ENNAVIA-A and ENNAVIA-B datasets.

### 2.3 Amino acid composition analysis

The amino acid composition of the experimentally verified antiviral peptides was analysed and compared to that of the experimentally verified non-antiviral peptide sequences and the random non-secretory peptide sequences extracted from UniProt. The composition analysis includes the peptides’ full sequences, the 10 N-terminal residues, and the C-terminal 10 residues.

### 2.4 Residue position preference analysis

Enrichment depletion logos (EDLogo)^[48]^ were created for the antiviral peptides’ sequences to identify any position-specific amino acid preferences that may exist. The experimentally validated non-antiviral peptide sequences were used as the baseline in the construction of the logo plots.

### 2.5 Features extraction

A variety of features was extracted from the peptides’ primary sequences. These features can be divided into two subcategories, amino acid-based descriptors and physicochemical descriptors. Only features that were non-zero for at least 20 samples were retained in the final feature vector.

#### 2.5.1 Composition descriptors

The peptides’ compositional descriptors were calculated based on the peptides’ amino acid, dipeptide, and tripeptide compositions for the conventional 20-amino acid alphabet. Additionally, descriptors were also calculated based on the reduced amino acid alphabets of Veltri et al.^[49]^, Thomas and Dill^[50]^, and the conjoint alphabet^[51]^. *g*-gap dipeptide and tripeptide compositions were calculated to account for the three-dimensional structure of the peptides^[52]^, with the values of the parameter *g* being 1, 2 and 3 for the dipeptide compositions, and 3 and 4 for the tripeptide compositions. Furthermore, conjoint triad, composition, transition and distribution^[53]^ and pseudo amino acid composition^[54]^ descriptors were also calculated.

#### 2.5.2 Physicochemical descriptors

The modlAMP package was employed for the calculation of global physicochemical descriptors and amino acid scale-based descriptors^[55]^. Global physicochemical features include molecular formula, sequence length, molecular weight, sequence charge, charge density, isoelectric point, instability index, aliphatic index^[56]^, aromaticity index^[57]^, hydrophobic ratio and the Boman index^[58]^. Amino acid scale-based descriptors include hydrophobicity^[59–63]^, side-chain bulkiness^[64]^, refractivity^[65]^, side-chain flexibility^[66]^, *α*-helix propensity^[67]^, transmembrane propensity^[68]^, polarity^[64,69]^, amino acid charges, AASI^[70]^, ABHPRK^[55]^, COUGAR^[55]^, Ez^[71]^, ISAECI^[72]^, MSS^[73]^, MSW^[74]^, PPCALI^[75]^, t_scale^[76]^, z3 ^[77]^, z5 ^[78]^ and pepArc^[55]^.

Additional physicochemical features were calculated based on amino acid properties detailed in the AAindex^[79]^. The peptides’ hydrophobicities were quantified using the amino acids’ hydrophobicities^[80,81]^, hydropathies^[82]^, retention coefficients in HPLC^[83]^ and partition energies^[84,85]^. Similarly, the peptide sequences’ hydrophilicities were characterised using descriptors based on the amino acid hydrophilicity scale^[86]^, the amino acids’ net charges^[87]^, polar requirements^[88]^ and fractions of site occupied by water^[89]^. Descriptors pertaining to sterics were obtained from the residues’ steric hindrance^[90]^ and bulkiness^[64]^ properties, while secondary structure features were calculated based on helical^[91]^ propensities. Furthermore, descriptors were also calculated from the side-chain interaction parameters^[92]^ and membrane-buried preference parameters^[93]^.

### 2.6 Machine learning approaches

Unsupervised and supervised machine learning approaches are employed in the current study. The former includes principal component analysis (PCA)^[94]^ and t-distributed Stochastic Neighbour Embedding (t-SNE)^[95]^ for visualising the data. The latter includes support vector machine (SVM)^[96]^, random forest (RF)^[97]^, and dense fully connected neural networks^[98]^ for creating supervised classifiers. The scikit-learn Python module is used for its PCA, t-SNE, SVM and RF implementations^[99]^.

SVMs were trialled using both a linear and non-linear radial base function (RBF) kernel. A grid search was employed for the tuning of the RF number of estimators, the maximum number of features, and the maximum depth hyperparameters, and the SVM regularization parameter C and kernel width parameter *γ*.

### 2.7 Neural network architecture and implementation

The Keras deep learning framework with a Tensorflow backend was used to build and train the deep-fully connected neural networks^[100]^.

The neural network’s input features are scaled to have minimum and maximum values of 0 and 1, respectively.

The optimal combination of neural network architecture and hyperparameters was selected using a randomized grid search strategy.

The first hidden layer has 1024 nodes, and is followed by two layers of 256 nodes each. Batch normalization^[101]^ is applied before the ReLU activation function for each hidden layer. To prevent overfitting to the training data, each hidden layer is followed by a Dropout regularization layer, with a rate of 0.30 ^[102]^. The output layer is a single node activated by the sigmoid function. As is common in binary classification neural networks, the binary cross-entropy loss function is employed.

It is defined as:

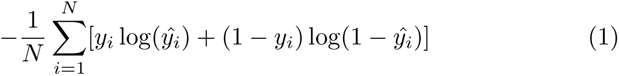

where *y_i_* is the true value of the *i^th^* sample, and *ŷ_i_* is the predicted value of the *i^th^* sample. As the predicted labels of all training data approach their respective true values, the value of the function approaches zero.

The optimal optimizer was found to be Adaptive Momentum (Adam), with an optimal initial learning rate of 0.05 and a decay of 0.0001. Adam utilises the following formula to update the neural network weights^[103]^:

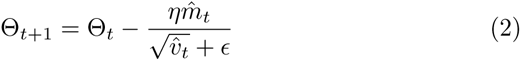

where the 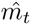 and 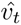 are the bias-corrected estimates of the mean and the variance of the gradients, respectively.

The neural networks were trained for 600 epochs, without stopping criteria. The model with the highest validation accuracy encountered during training was retained for each of the cross-validation splits.

### 2.8 Transfer learning

As the dataset of peptides with anti-coronavirus is small, numbering only 109 peptides, transfer learning was used to train the models for the ENNAVIA-C and ENNAVIA-D datasets. Models originally trained for each cross-validation fold for ENNAVIA-A and ENNAVIA-B, respectively, were used to initialize the weights for the neural network models of the corresponding cross-validation folds for ENNAVIA-C and ENNAVIA-D, respectively. The neural network models were then trained for 600 epochs, without stopping criteria. The model with the highest validation accuracy encountered during training was retained for each of the cross-validation splits.

### 2.9 Performance evaluation

A number of standard metrics are employed for the evaluation of the presented models’ performance, specifically accuracy (Acc), sensitivity (Sn), specificity (Sp), the Matthews correlation coefficient (MCC), and the receiver operating characteristic (ROC) curve. Confidence intervals are provided at the 95% level of significance.

The first four metrics are defined by the following equations:

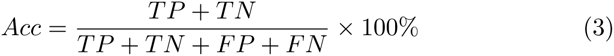

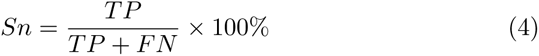

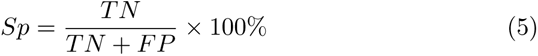

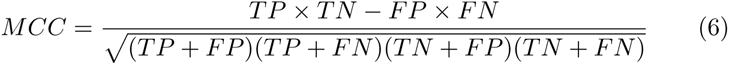

where

- TP = True positives: the number of correctly predicted positive (antiviral) peptides.
- FP = False positives: the number of non-antiviral peptides incorrectly predicted as being antiviral.
- TN = True negatives: the number of correctly predicted negative (non-antiviral) peptides.
- FN = False negatives: the number of anticancer peptides incorrectly predicted as being non-antiviral.

## 3 Results

The dataset of peptide sequences was subjected to an amino acid composition analysis and residue position preference analysis. Feature vectors comprising the peptides’ physicochemical descriptors, compositional descriptors, and all descriptors were constructed and visualised in two-dimensional space using principal component analysis (PCA) and t-stochastic neighbour embedding (t-SNE) plots. Plots created using both methods show an incomplete separation of the positive and negative classes. Finally, three machine learning classifiers, namely support vector machines, random forests and neural networks, are trained on the dataset’s feature vectors, and the antiviral activity prediction results are evaluated.

### 3.1 Amino acid composition analysis

To identify if particular amino acid residues are more prevalent in antiviral and anti-coronavirus peptides, an amino acid residue composition analysis was performed. The amino acid compositions of anti-coronavirus peptides, antiviral peptides, experimentally validated non-antiviral peptides, and random non-antiviral peptide sequences are illustrated in Figure 1.

**Figure 1:**
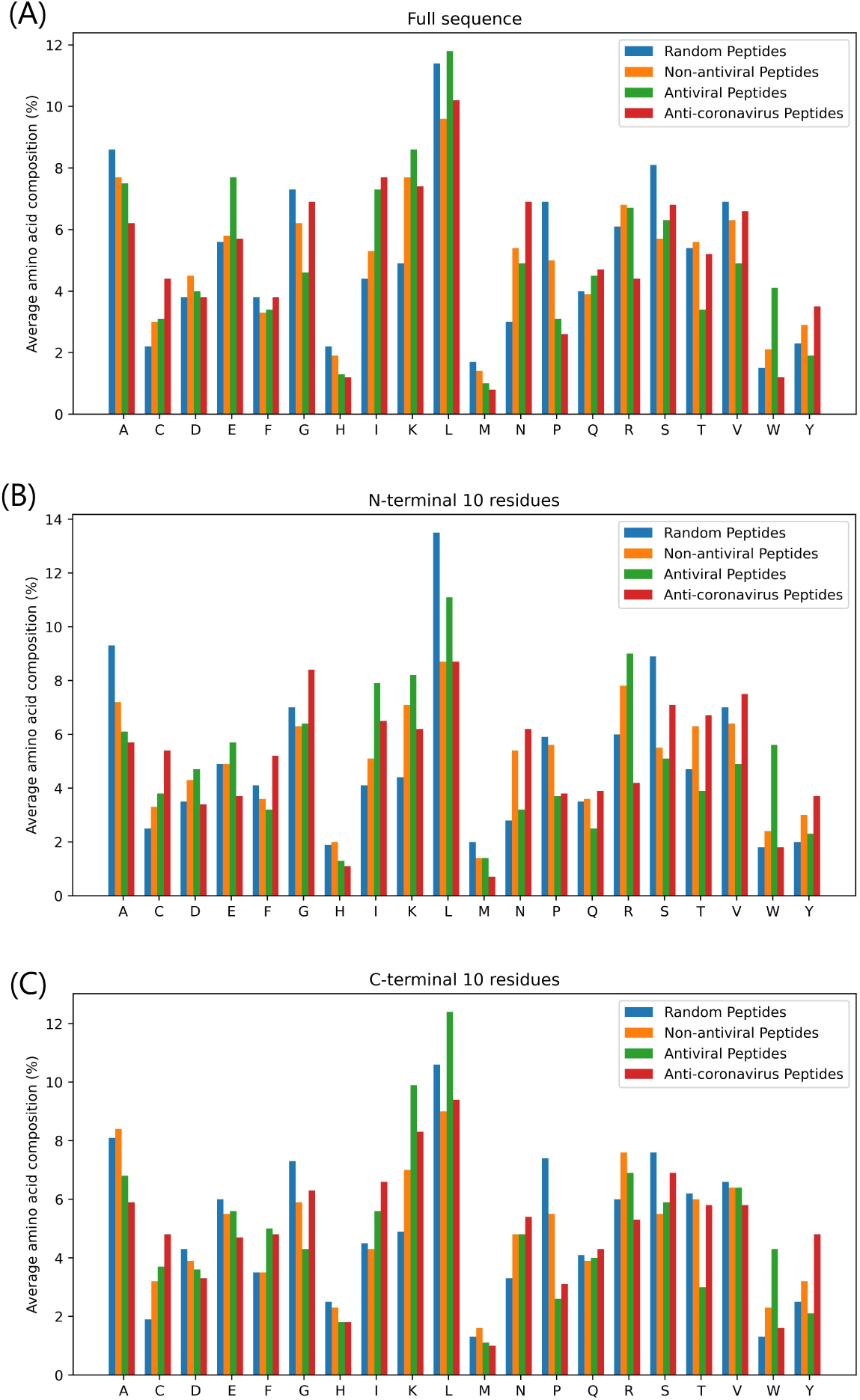
Percentage average amino acid residue composition of the (A) full sequences, (B) N-terminal 10 residues, and (C) C-terminal 10 residues of anti-coronavirus peptides (red), experimentally validated antiviral peptides (green), experimentally validated non-antiviral peptide (orange) and non-antiviral peptides randomly extracted from UniProt proteins (blue). One-letter amino acid codes are given for the residues on the x-axis.

Interestingly, antiviral and anti-coronavirus peptides are enriched in the cysteine and the hydrophobic isoleucine residue, and depleted in proline and histidine. While antiviral peptides in general exhibit enrichment in lysine and tryptophan, this is not observed for the specifically anti-coronavirus peptides. Similarly, antiviral peptides are depleted in glycine and valine, while anti-coronavirus peptides are enriched in these residues. While the amino acid composition for anti-coronavirus peptides is based on a limited sample size, it does suggest that the composition requirements for peptides to possess activity against coronaviruses differ from the composition requirements for activity against viruses in general.

### 3.2 Residue position preference analysis

The possibility of a preference existing for certain amino acid residues at certain positions in the peptides’ primary sequence, an enrichment-depletion logo plot was produced (Figure 2) for the experimentally validated antiviral peptides. The experimentally validated non-antiviral peptides were used to establish a baseline for the plot.

**Figure 2:**
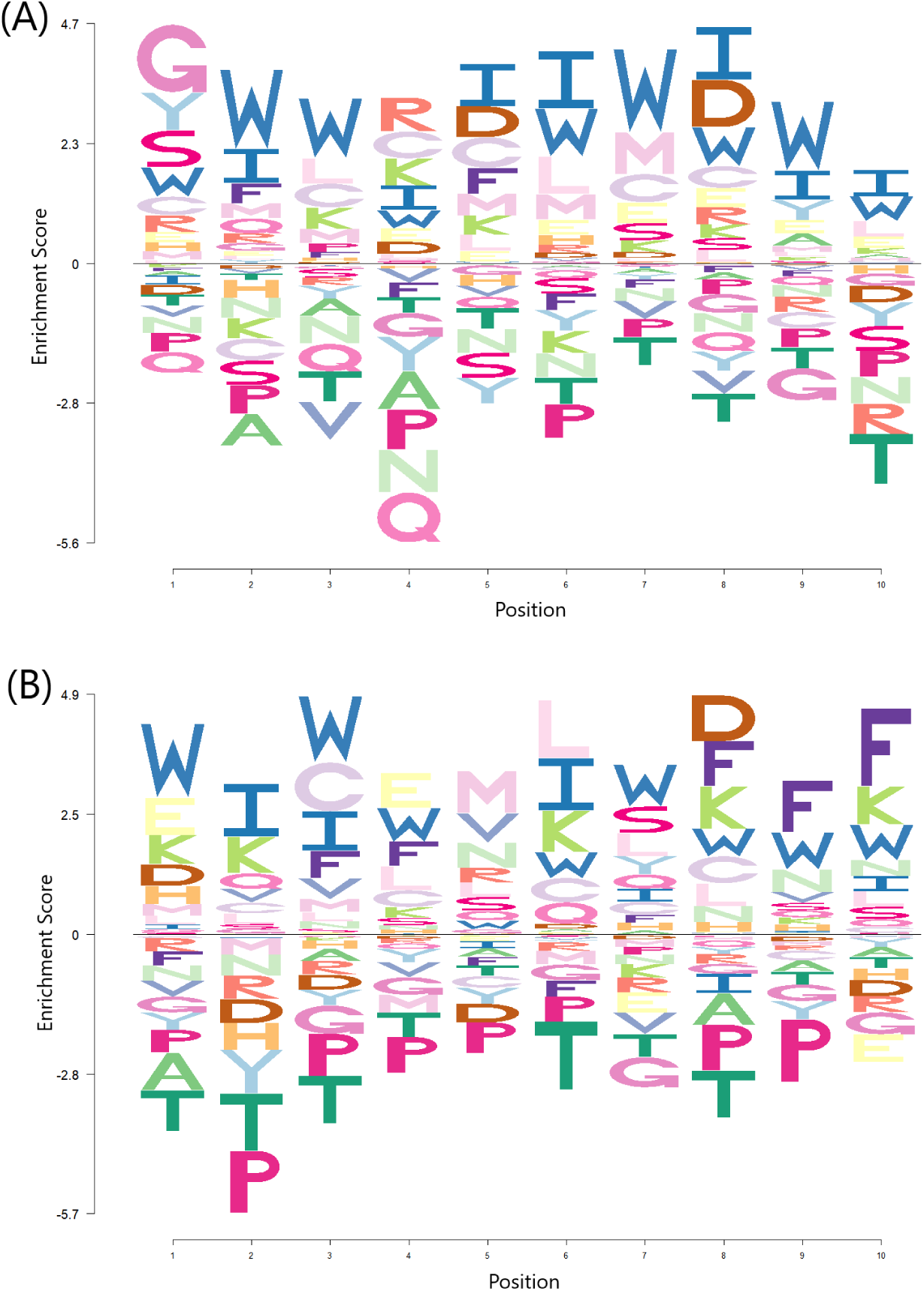
Enrichment-depletion logo plot of (A) N-terminal 10 residues and (B) C-terminal 10 residues of experimentally validated antiviral peptides of the ENNAVIA-A dataset. Data is scaled to account for the background probability of each amino acid, based on the experimentally validated non-antiviral peptides dataset.

The first inspection of the logo plot suggests that antiviral peptides are enriched in tryptophan at most positions. This is consistent with the afore-mentioned amino acid composition analysis. More specifically, however, antiviral peptides appear to be enriched in glycine at position 1, and have a preference for a positively charged residue at position 4. Conversely, they are enriched in aspartic acid at position 5 and 8, and the third-last residue. Enrichment is also observed in phenylalanine at the three C-terminal positions. Again, in agreement with the amino acid composition analysis, antiviral peptides are depleted in proline and tryptophan at all positions.

### 3.3 Data Visualisation

#### 3.3.1 Principal Component Analysis (PCA)

Principal component analysis (PCA) was carried out on the ENNAVIA-A dataset for all computed descriptors, only the physicochemical descriptors, and only the compositional descriptors subsets (Figure 3). While a separation does exist between the experimentally verified antiviral and non-antiviral peptides, it is incomplete, and the two classes are significantly overlapped.

**Figure 3:**
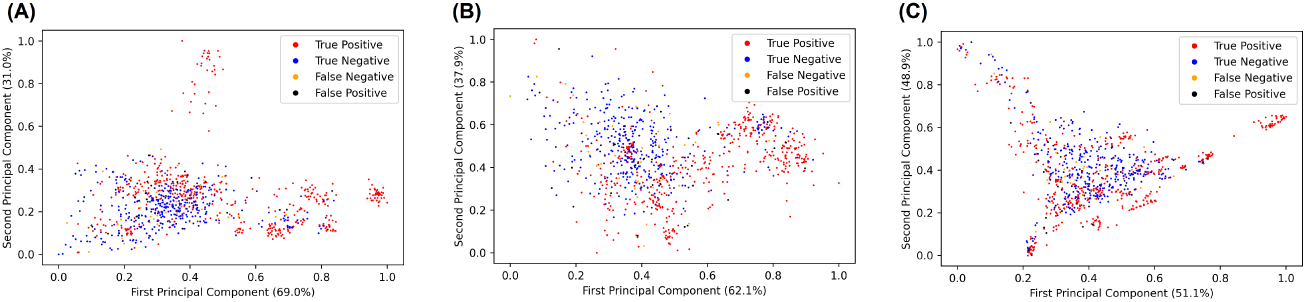
Principal component analysis of (A) all descriptors, (B) only the physicochemical descriptors and (C) only the compositional descriptors. Experimentally validated antiviral peptides (positives) are coloured red, experimentally validated non-antiviral peptides (negatives) are coloured yellow, false-positives are coloured black, false-negatives are coloured blue.

#### 3.3.2 T-Distributed Stochastic Neighbour Embedding (t-SNE)

To complement the PCA analysis, a t-distributed Stochastic Neighbour Embedding (t-SNE) analysis was conducted for the experimentally verified antiviral and non-antiviral peptides, again for all computed descriptors, only the physicochemical descriptors, and only the compositional descriptors subsets (Figure 4). As with the results of the PCA analysis, the interclass separation is incomplete, although it is clearly greater.

**Figure 4:**
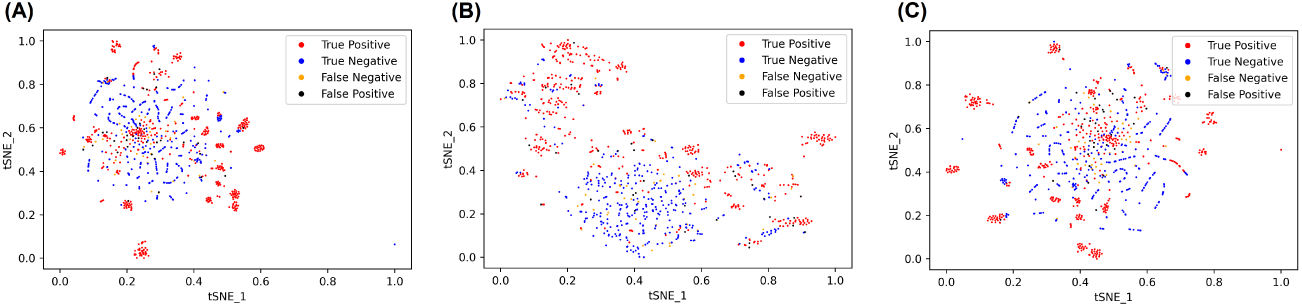
t-SNE visualisation of (A) all descriptors, (B) only the physicochemical descriptors and (C) only the compositional descriptors. Experimentally validated antiviral peptides (positives) are coloured red, experimentally validated non-antiviral peptides (negatives) are coloured yellow, false-positives are coloured black, false-negatives are coloured blue.

### 3.4 Antiviral activity prediction

The principal aim of this study was to train and evaluate a selection of machine learning classifiers for the prediction of peptide antiviral activity.

Tenfold cross-validation was employed for the evaluation of the classifiers’ robustness and predictive power. Additionally, the ten models trained for each classifier under tenfold cross-validation were ensembled and further evaluated through the use of the external, independent test set. The accuracy, Matthews correlation coefficient, sensitivity and specificity parameters, together with their respective confidence intervals, are reported for each model. Receiver operating characteristic curves (ROC) with calculated area under the curve (AUC) values are also given for both final neural network models. The support vector machine (SVM), random forest (RF) and neural network (NN) performance metrics are tabulated in Table 1.

**Table 1:**
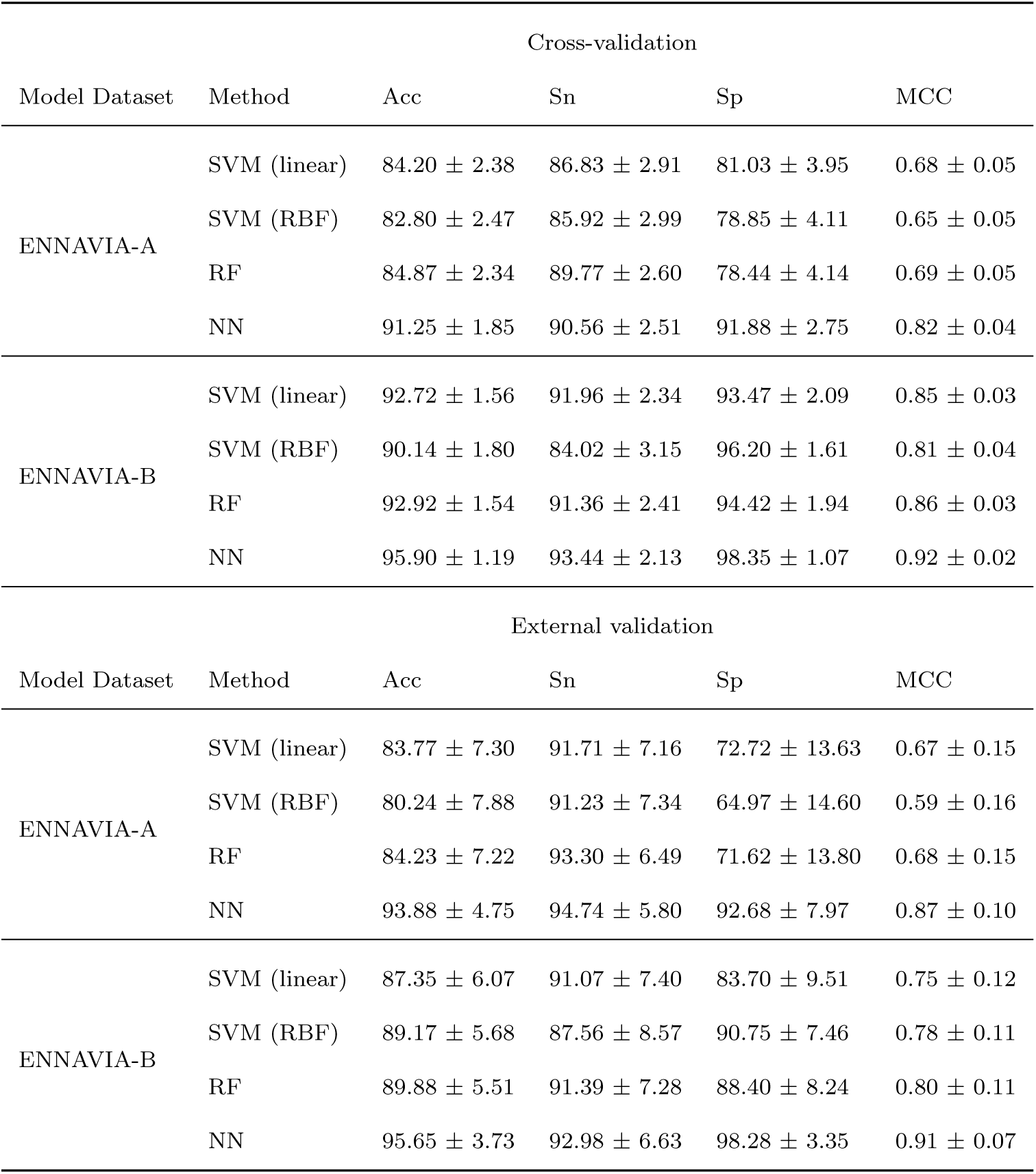
Cross-validation and external validation statistical parameters for SVM with a linear kernel, SVM with a RBF kernel, RF and NN models trained on the ENNAVIA-A and ENNAVIA-B datasets

| Model Dataset | Method | Cross-validation |  |  |  |
| --- | --- | --- | --- | --- | --- |
|  |  | Acc | Sn | Sp | MCC |
| ENNAVIA-A | SVM (linear) | $84.20 \pm 2.38$ | $86.83 \pm 2.91$ | $81.03 \pm 3.95$ | $0.68 \pm 0.05$ |
| | SVM (RBF) | $82.80 \pm 2.47$ | $85.92 \pm 2.99$ | $78.85 \pm 4.11$ | $0.65 \pm 0.05$ |
| | RF | $84.87 \pm 2.34$ | $89.77 \pm 2.60$ | $78.44 \pm 4.14$ | $0.69 \pm 0.05$ |
| | NN | $91.25 \pm 1.85$ | $90.56 \pm 2.51$ | $91.88 \pm 2.75$ | $0.82 \pm 0.04$ |
| ENNAVIA-B | SVM (linear) | $92.72 \pm 1.56$ | $91.96 \pm 2.34$ | $93.47 \pm 2.09$ | $0.85 \pm 0.03$ |
| | SVM (RBF) | $90.14 \pm 1.80$ | $84.02 \pm 3.15$ | $96.20 \pm 1.61$ | $0.81 \pm 0.04$ |
| | RF | $92.92 \pm 1.54$ | $91.36 \pm 2.41$ | $94.42 \pm 1.94$ | $0.86 \pm 0.03$ |
| | NN | $95.90 \pm 1.19$ | $93.44 \pm 2.13$ | $98.35 \pm 1.07$ | $0.92 \pm 0.02$ |
| Model Dataset | Method | External validation |  |  |  |
|  |  | Acc | Sn | Sp | MCC |
| ENNAVIA-A | SVM (linear) | $83.77 \pm 7.30$ | $91.71 \pm 7.16$ | $72.72 \pm 13.63$ | $0.67 \pm 0.15$ |
| | SVM (RBF) | $80.24 \pm 7.88$ | $91.23 \pm 7.34$ | $64.97 \pm 14.60$ | $0.59 \pm 0.16$ |
| | RF | $84.23 \pm 7.22$ | $93.30 \pm 6.49$ | $71.62 \pm 13.80$ | $0.68 \pm 0.15$ |
| | NN | $93.88 \pm 4.75$ | $94.74 \pm 5.80$ | $92.68 \pm 7.97$ | $0.87 \pm 0.10$ |
| ENNAVIA-B | SVM (linear) | $87.35 \pm 6.07$ | $91.07 \pm 7.40$ | $83.70 \pm 9.51$ | $0.75 \pm 0.12$ |
| | SVM (RBF) | $89.17 \pm 5.68$ | $87.56 \pm 8.57$ | $90.75 \pm 7.46$ | $0.78 \pm 0.11$ |
| | RF | $89.88 \pm 5.51$ | $91.39 \pm 7.28$ | $88.40 \pm 8.24$ | $0.80 \pm 0.11$ |
| | NN | $95.65 \pm 3.73$ | $92.98 \pm 6.63$ | $98.28 \pm 3.35$ | $0.91 \pm 0.07$ |

A grid search strategy was employed for the optimization of SVM and RF hyperparameters.

The SVM classifier achieved its best performance with the regularization parameters *C* = 1 and *C* = 10 for the linear and RBF kernels, respectively, and the kernel coefficient *γ* = 1.5 × 10^−4^ for the non-linear kernel. The SVM classifiers, both with a linear and non-linear kernel, perform worse than the RF and NN approaches, with cross-validation accuracies of 84.2% and 82.8%, and MCCs of 0.68 and 0.65, respectively on the ENNAVIA-A dataset.

The optimal RF hyperparameters differed depending on the dataset used. For the ENNAVIA-A dataset, optimal performance was observed with 124 estimators, a maximum tree depth of 10 and a maximum of 80 features, achieving a cross-validation accuracy and MCC of 84.9% and 0.69, respectively. For the ENNAVIA-B dataset, meanwhile, optimal performance was observed with 512 estimators, unrestricted tree depth and a maximum of 13 features.

The neural network approach, however, achieves the best predictive performance of all machine learning approaches trialled, with an accuracy and MCC scores of 93.88% and 0.87 on the ENNAVIA-A external test set, and 95.65% and 0.91 on the ENNAVIA-B external test set. Furthermore, the neural network achieves a very good balance between sensitivity and specificity, 94.74% and 92.68% for ENNAVIA-A. Receiver operating characteristic (ROC) (Figure 5) curves were produced to further evaluate the neural networks’ robustness, as were the corresponding area under the curve (AUC) values, which were calculated as 0.93 and 0.98 for the ENNAVIA-A and ENNAVIA-B models, respectively.

**Figure 5:**
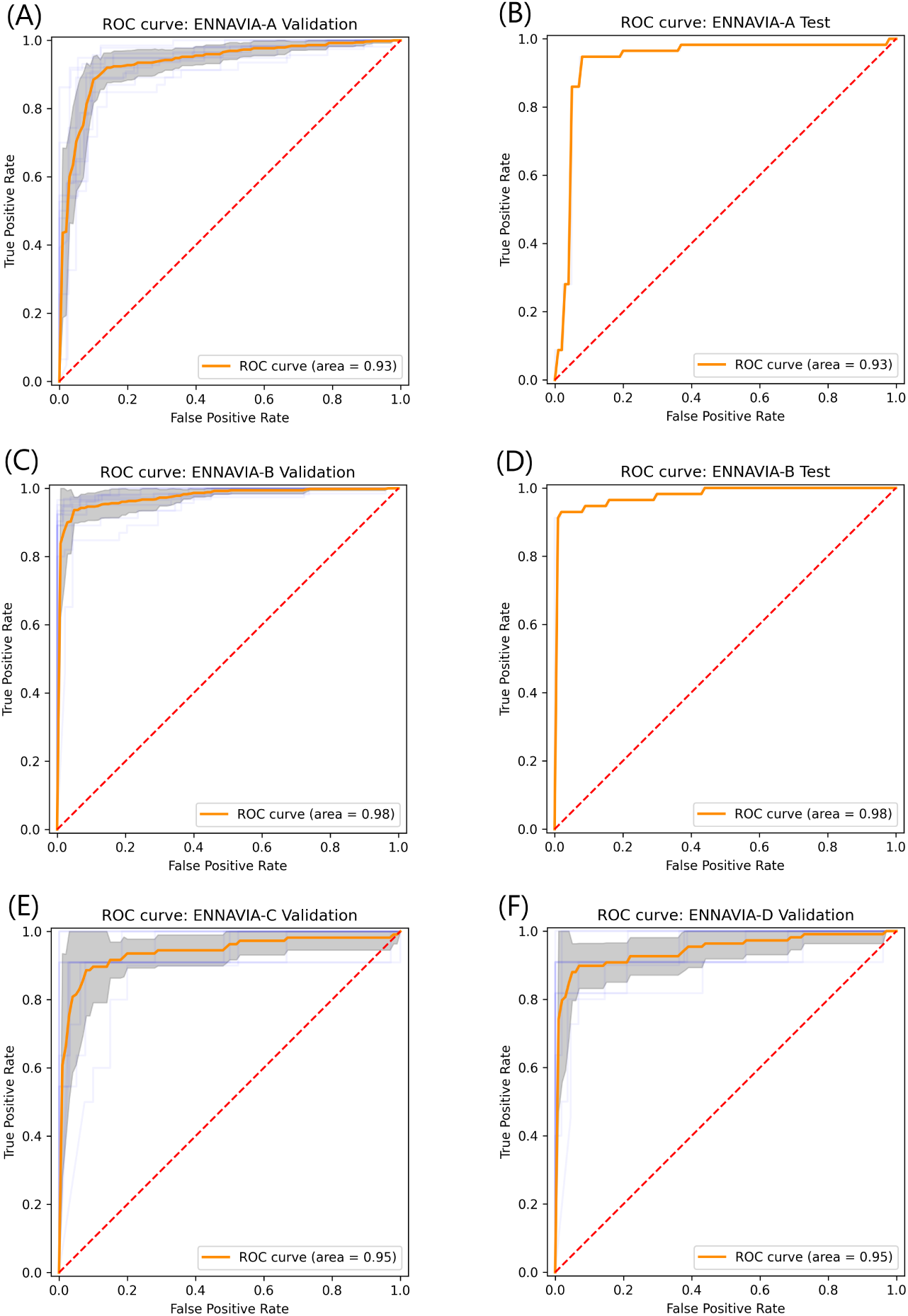
Receiver operating characteristic (ROC) plots and associated Area Under the Curve (AUC) values for model performance on (A,B) ENNAVIA-A tenfold cross-validation set and external test set (C,D) ENNAVIA-B tenfold cross-validation set and external test set and (E,F) ENNAVIA-C and ENNAVIA-D tenfold cross-validation sets. The ENNAVIA-A model achieves an AUC value of 0.93, ENNAVIA-B achieves and AUC value of 0.98, and ENNAVIA-C and ENNAVIA-D both achieve values of 0.95.

As the neural networks’ performance was superior to the SVM and RF approaches, it was deemed as the best model for the prediction of peptide antiviral activity and further studied.

### 3.5 Comparison with existing peptide antiviral activity prediction methods

To establish the utility of ENNAVIA in the context of prediction methods already described in the literature, ENNAVIA was benchmarked against three existing antiviral peptide prediction methods, specifically AVPpred^[31]^, the method of Chang et al.^[35]^, AntiVPP^[32]^, Meta-iAVP^[104]^ and FIRMAVP^[34]^. Detailed results are given in Table 2.

**Table 2:**
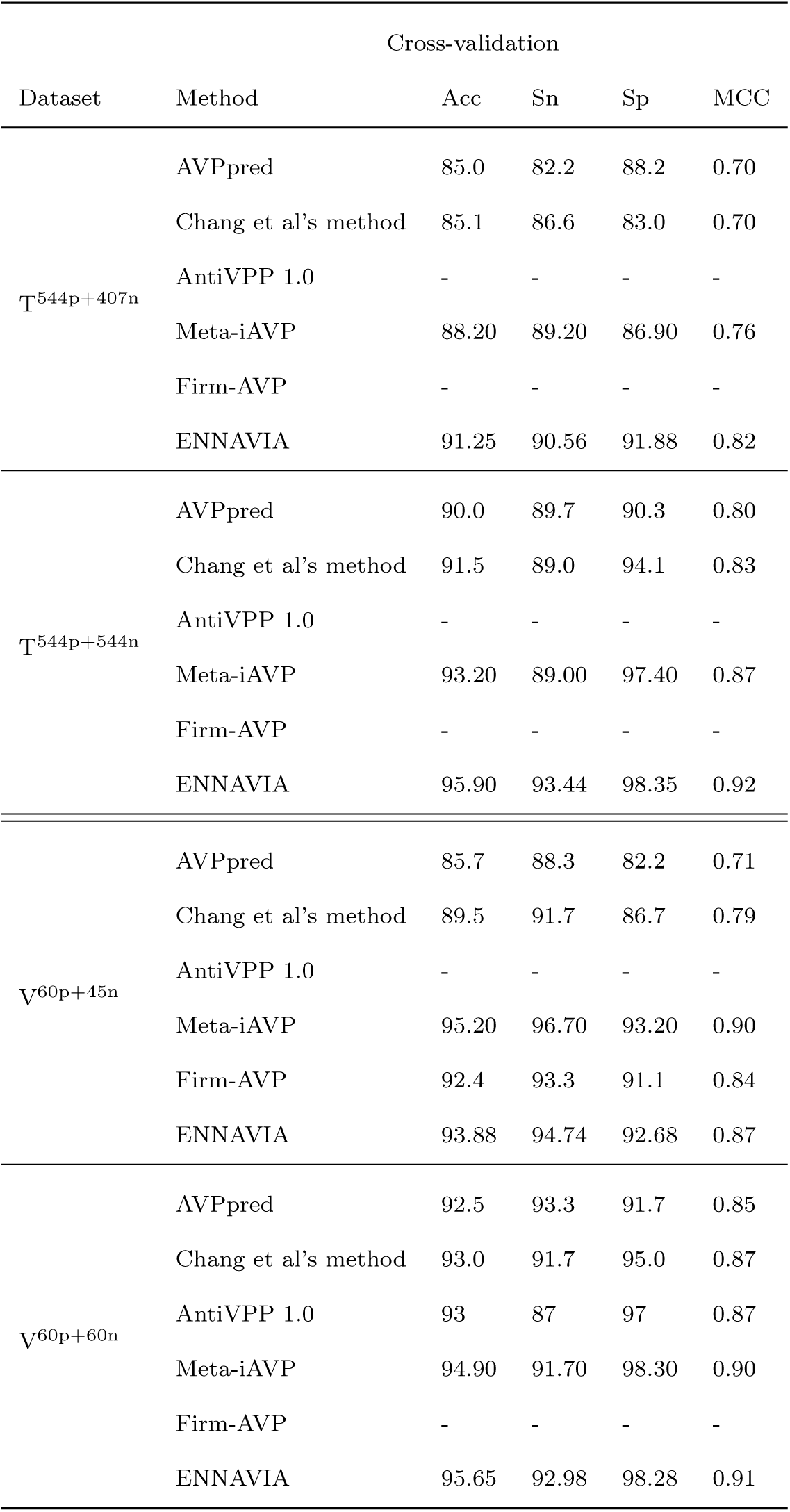
Cross-validation and external validation performance comparison between ENNAVIA and existing methods for the prediction of peptide antiviral activity.

The results presented in Table 2 are reproduced from the respective articles describing the methods. It must be noted, however, that the results for Meta-iAVP and AntiVPP 1.0 could not be reproduced. Independent evaluation of the Meta-iAVP via its webserver on the V^60p+45n^ dataset resulted in Acc, Sn and Sp values of 81.0%, 83.3% and 77.8%, respectively. Similarly, evaluation of the AntiVPP 1.0 software on the V^V60p+60n^ dataset resulted in Acc, Sn and Sp values of 81.6%, 76.6% and 86.6%, respectively. Contact with the corresponding authors of these articles was attempted prior to publication, however, we have not received a response to our queries prior to publication.

### 3.6 Anti-coronavirus activity prediction

A recent study by Pang et al. described a machine learning method for the identification of anti-coronavirus peptides through imbalanced learning strategies^[37]^. This study utilises the datasets created by Pang et al. and employs transfer learning to adapt the ENNAVIA-A and ENNAVIA-B models to the task of anti-coronavirus peptide prediction. For both ENNAVIA-A and ENNAVIA-B, the neural network weights of each of the ten models trained under cross-validation are transferred to their corresponding models for anti-coronavirus peptide prediction, which are then trained on their respective datasets. The accuracy, Matthews correlation coefficient, sensitivity and specificity parameters, together with their respective confidence intervals, are reported for each model. Receiver operating characteristic curves (ROC) with the calculated area under the curve (AUC) values are also given for both final neural network models (Fig 5). The anti-coronavirus peptide prediction performance obtained by each model is compared with the results obtained by Pang et al. As the size of the anti-coronavirus peptide dataset is extremely limited, and neural network performance typically increases with the amount of data available, validation is limited to ten-fold cross-validation. Detailed results are given in Table 3.

**Table 3:** Performance evaluation of ENNAVIA in prediction of anticoronavirus peptides, and comparison with the methods of Pang et al.

| Dataset | Method | Acc | Sn | Sp | MCC |
| --- | --- | --- | --- | --- | --- |
| Anti-CoV vs. Non-AVP | Pang et al’s method | 85.32 | 85.71 | 85.31 | 0.305 |
| ENNAVIA-C | ENNAVIA | $94.95 \pm 1.99$ | $91.64 \pm 5.20$ | $95.96 \pm 2.04$ | $0.87 \pm 0.05$ |
| Anti-CoV vs. Random | Pang et al’s method | 97.72 | 100 | 97.66 | 0.730 |
| ENNAVIA-D | ENNAVIA | $97.29 \pm 1.25$ | $89.82 \pm 5.68$ | $98.77 \pm 0.93$ | $0.91 \pm 0.03$ |

### 3.7 Descriptor-set specific results

To ascertain the extent to which a given set of features can contribute to the correct prediction of peptide antiviral activity, neural networks were trained on subsets of the feature space. The validation results obtained by these neural networks trained on the peptides’ physicochemical features, dipeptide composition, dipeptide *g*-gap composition and tripeptide composition are detailed in Table 4.

**Table 4:**
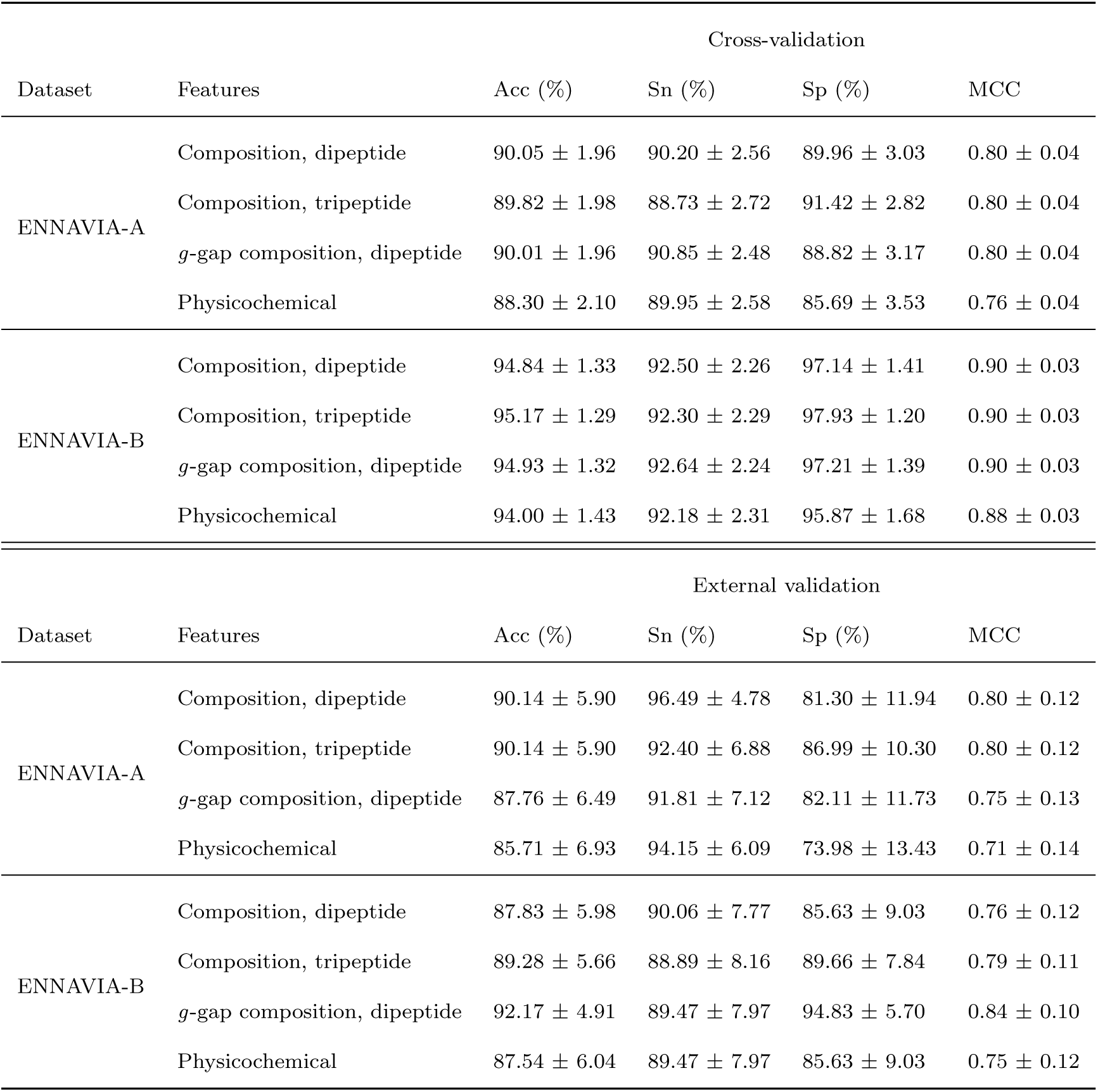
Validation statistics achieved by neural network models trained on subsets of the feature space. The *g*-gap parameter *g* =1,2,3.

None of the reduced subset models trained achieve performance better than the hybrid model trained on both compositional and physicochemical descriptors, validating the choice of the hybrid model as the principal approach.

#### 3.7.1 Dipeptide and tripeptide composition

Information about local sequence order can be relayed to a machine learning method through the use of dipeptide and tripeptide composition descriptors. A peptide’s dipeptide and tripeptide composition can be defined as the percentage of a given dipeptide or tripeptide in the sequence. These features also have the added benefit of capturing the peptide’s chemical nature. Both the dipeptide-based model and the tripeptide-based model achieve good results, with accuracies of 90.1% and 89.8%, respectively, and MCC values of 0.80.

#### 3.7.2 *g*-gap composition

*g*-gap compositions, defined as the proportion of a pair of amino acids separated by 1, 2 or 3 residues, are a useful descriptor as they correspond to residues that may be proximate to one another in three-dimensional space. As peptides often possess secondary structure upon interaction with their targets, this information allows for the capturing of the chemical environment that the peptide presents to its target. Models trained on *g*-gap dipeptide composition do not perform better than those trained on conventional dipeptide composition, achieving an accuracy and MCC of 90.0% and 0.80.

#### 3.7.3 Physicochemical

Models trained on physicochemical features, such as charge, amphiphilicity and charge, achieve an accuracy and MCC of 88.3% and 0.76, respectively.

Although this performance is poorer than that achieved by the models trained on compositional features, it is only marginally so, and still demonstrates predictive capability.

## 4 Discussion

The need for novel anti-viral drugs, especially in the context of the COVID-19 pandemic, is great. Interest in the development of novel peptide-based therapeutics has increased in recent years, even as the number of new drugs approved each year declines and the cost of drug research and development grows. More specifically, antiviral peptides represent a promising class of novel drug candidates. Despite extensive research having been conducted on the relationship between the conformations of various bioactive peptides and their biological activities^[105–107]^, understanding of this relationship remains insufficient the accurate de-novo design of novel peptide drugs, especially antiviral peptide drugs, which compared to antimicrobial peptides are less numerous in the literature, and consequently less studied. Molecular dynamics simulations can reveal insights into activity, but are time-consuming and largely unsuitable for bulk-screening of peptide sequences.

An accurate computational method for the prediction of peptide antiviral activity from the primary sequence alone would facilitate a more rapid exploration of the peptide chemical space, and lower the cost of research and development by reducing the need for chemical synthesis and laboratory evaluation of peptide antiviral activity. With a view to accelerating the screening and design of new antiviral peptide drugs, the present study focuses on the combination of compositional and physicochemical descriptors with a deep neural network architecture to create an *in silico* method for a more accurate classification of peptides as either antiviral or non-antiviral, and additionally the prediction of peptide anti-coronavirus activity specifically, solely on the basis of their primary sequence.

To facilitate as direct a comparison as possible with existing antiviral peptide prediction methods, the dataset of Thakur et al.^[31]^ was adapted for use in this study. Peptide sequences comprising non-natural amino acids or with a length outside the 7-40 amino acid range were excluded. 577 of the original 604 antiviral peptides remain in the ENNAVIA datasets. Two negative datasets are used in this study: the ENNAVIA-A dataset includes 420 experimentally evaluated non-antiviral peptides, while the ENNAVIA-B dataset includes 597 random peptide sequences as the negative samples.

Compositional and physicochemical descriptors were employed for the construction of feature vectors from the peptides’ primary sequences, and a selection of machine learning methods were evaluated for the peptide antiviral activity prediction task through both tenfold cross-validation and validation on an external test set. Deep neural networks proved most promising, and their architecture was, therefore, further optimised and evaluated.

The neural network model with five hidden layers was found to achieve optimal performance. On the ENNAVIA-A dataset, a tenfold cross-validated accuracy, sensitivity and specificity of 91.3%, 90.6% and 91.9%, was achieved, clearly demonstrating that the neural network model is capable of accurately identifying antiviral peptides among non-antiviral peptides. ENNAVIA’s predictive performance was compared to existing methods, especially the existent state-of-the-art, Meta-iAVP, which exhibited a cross-validated accuracy, sensitivity and specificity of 88.2%, 89.2% and 86.9%, respectively, on the T^504p+407n^ dataset. ENNAVIA’s performance surpasses that of Meta-iAVP and other existent models on all metrics, designating it a new state-of-the-art model for antiviral peptide prediction.

Similarly, neural network models were trained and evaluated on the ENNAVIA-B dataset, achieving cross-validated accuracy, sensitivity and specificity of 95.9%, 93.4% and 98.6%, respectively, demonstrating that ENNAVIA can distinguish between antiviral peptides and random peptide sequences. A comparison of performance on this dataset to existing methods again establishes ENNAVIA as the best-in-class method for antiviral peptide prediction, surpassing the previously best accuracy, sensitivity and specificity of 93.2%, 89.0% and 97.4% achieved by meta-iAVP on the _T_504p+504n _dataset._

Recently, Pang et al. published a study that employed random forests with imbalanced learning strategies for the identification of anti-coronavirus peptides. Notably, the anti-coronavirus peptide dataset is small, with only a total of 139 peptide sequences. Despite the small number of positive samples available for training, respectable validation statistics were achieved, with a sensitivity, specificity and MCC of 85.7%, 85.3% and 0.31 with non-antivirus peptides as the negative dataset, and 100%, 97.7% and 0.73 with random peptide sequences as the negative dataset.

To expand the scope of the current study to include the facilitation of rapid screening of peptides for anti-coronavirus activity specifically, two additional datasets which include the anti-coronavirus peptides from the dataset of Pang et al. as the positive samples were constructed: ENNAVIA-C and ENNAVIA-D, which use the negative peptides from the ENNAVIA-A and ENNAVIA-B datasets, respectively. As the number of positive samples is too small to accurately train neural network models, transfer learning was employed, whereby the already-trained weights of the ENNAVIA-A and ENNAVIA-B models were transferred to the ENNAVIA-C and ENNAVIA-D models, respectively, and further fine-tuned to the anti-coronavirus peptide prediction task. The ENNAVIA-C model achieved a sensitivity, specificity and MCC of 91.6%, 96.0% and 0.87, representing a significant improvement on the work of Pang et al. The ENNAVIA-D model, similarly, achieved good performance, with a sensitivity, specificity and MCC of 89.8%, 98.8% and 0.91, respectively, out-performing the method of Pang et al. in specificity, although not sensitivity.

Nonetheless, as neural networks’ predictive power scales with the quantity of data available for training, further improvements in predictive performance for both the antiviral and anti-coronavirus predictive models could be achieved as the literature on antiviral peptides expands, and the number of peptide sequences available for training increases.

To conclude, the limited quantity of available experimentally validated data and the incomplete understanding of the mechanism of peptide antiviral activity continue to pose challenges for the research community. In an effort to overcome these challenges, this study described ENNAVIA, a collection of novel *in silico* peptide antiviral and anti-coronavirus activity classifiers. The classifiers, which employ a deep neural network architecture and benefit from a rich feature-space, achieve predictive power that surpasses the state-of-the-art. This work complements a suite of existing *in silico* classifiers developed by the authors, which includes methods for the prediction of peptide anticancer and hemolytic activity, and peptide tertiary structure. The authors believe that the results of this work, in combination with the aforementioned methods, will enable better *in-silico* design of novel peptide-based antiviral and anti-coronavirus therapeutics, thereby reducing the cost and time required for the design phase, helping to drive medicinal chemistry into an unprecedented revolution.

## 5 Web server implementation

For the benefit of the scientific community, the ENNAVIA classifier is available as a user-friendly, publicly accessible web server online at https://research.timmons.eu/ennavia. The web server is capable of predicting peptides’ antiviral activity based on the primary sequence. Input peptide sequences are restricted to only the 20 20 natural amino acids; non-natural amino acids are not supported. The web server includes many features, and models trained on both the ENNAVIA-A (T^504p+406n^) and ENNAVIA-B dataset (T^504p+504n^) are available for prediction.

### 5.1 Peptide antiviral activity prediction

Peptide antiviral activities can be predicted for both a single sequence and a batch of sequences. Peptide sequences should be provided in the standard FASTA format. The maximum batch size is variable depending on the length of the sequences; longer sequences necessitate smaller batch sizes. The prediction will be carried out by the ensemble of trained neural networks, and the average score will be returned, which corresponds to the probability of the peptide sequence possessing antiviral activity. Probabilities are given on a scale of 0-1, whereby 0 and 1 are most probably non-antiviral, and most probably antiviral, respectively.

### 5.2 Mutation analysis

Mutation analysis may be carried out on single peptide sequences, by selecting the mutation analysis option and inputting the residue number to be mutated. Mutant sequences will be created by substituting the residue at the specified position with each of the other 20 natural amino acids. The probability of each of the mutant sequences possessing antiviral activity will be returned by the chosen neural network model.

### 5.3 Residue scan

Residue scans, such as, for instance, an alanine scan, are available for single peptide sequences, by choosing the residue scan option and selecting the amino acid residue to be scanned with. Mutant sequences are attained by substituting successive residues with the selected amino acid residue. The probability of the native and mutant sequences possessing antiviral activity will be returned by the selected neural network model.

## Data Availability

All data generated or analysed during this study are available for download at https://research.timmons.eu/ennavia

## Acknowledgements

The authors would also like to thank University College Dublin for the Research Scholarship granted to P.B.T.

